# Costs and benefits of toxin production in a dinoflagellate

**DOI:** 10.1101/2020.07.12.199380

**Authors:** Fredrik Ryderheim, Erik Selander, Thomas Kiørboe

## Abstract

Many phytoplankton respond to chemical cues from grazers by upregulating defensive capabilities. Inducible defences like these are often assumed to come at a cost to the organism, but these trade-offs have not been experimentally established. A reason for this may be that costs only become evident under resource limiting conditions. Here, we exposed the toxin-producing dinoflagellate *Alexandrium minutum* to chemical cues from copepods under different levels of nitrogen limitation. Induced cells had higher cellular toxin content and a larger fraction of the cells were rejected by a copepod, demonstrating the clear benefits of toxin production. Induced cells also had a higher carbon and nitrogen content, despite an up to 25% reduction in cell size. Unexpectedly, induced cells seemed to grow faster than controls, likely owing to a higher nutrient affinity due to reduced size. We thus found no clear trade-offs, rather the opposite. However, we argue that indirect ecological costs that do not manifest under laboratory conditions are important and that the induction of toxins specific to particular defences prevents the cells from constantly synthesizing the large array of secondary metabolites that they are capable of producing.

## Introduction

Dinoflagellates of the genus *Alexandrium* produce paralytic shellfish toxins (PSTs), intracellular secondary metabolites that may have defensive capabilities (Selander *et al.* 2006; Xu & Kiørboe 2018). Indeed, toxicity as a defence mechanism against grazers is the favoured explanation for the evolution of algal toxins (Turner & Tester 1997; Smetacek 2001; Xu & Kiørboe 2018).

Studies dedicated to defence mechanisms in phytoplankton have often focused on the benefits of the defence, but have rarely established potential costs (Pančić & Kiørboe 2018). So far, many experimental assessments have suggested toxin production trade-offs to be insignificant. Thus, the growth rate of toxic and non-toxic strains of the same species, or the growth of grazer-induced versus non-induced cells of the same strain but with very different toxin contents appear to be identical (John & Flynn 2000; Selander *et al.* 2006, 2008). Blossom *et al.* (2019) compared several species and strains of *Alexandrium* spp. and similarly did not find any correlation between growth rate and toxin production under light replete conditions, and even a positive correlation under limiting light. Brown & Kubanek (2020) demonstrated a negative relation between toxin content and growth rate in *Alexandrium minutum* exposed to lysed cells of various other species of dinoflagellates, thus suggesting a trade-off. However, the correlation may well have been spurious and driven independently by allelochemical substances in the lysed cells (Windust *et al.* 1996). It is well documented that many dinoflagellates produce such dissolved allelochemicals that reduce the growth rate of other cells (Legrand *et al.* 2003). Significant costs of predator-induced toxin production have so far only been convincingly demonstrated in diatoms that produce domoic acid, but here the defence mechanisms are unclear (Lundholm *et al.* 2018). However, if defence mechanisms are adaptive there must be associated costs; otherwise, non-defended species or strains would be outcompeted and all species would be equally defended. Also, toxin production is inducible; i.e., it is upregulated in the presence of grazers cues, as seen in *Alexandrium* spp. dinoflagellates (e.g. Selander *et al.* 2006; Bergkvist *et al.* 2008; Griffin *et al.* 2019) and some toxic strains of the diatom *Pseudo-nitzschia* (Lundholm *et al.* 2018; Selander *et al.* 2019). According to optimal defence theory, inducible defences are favoured when risks vary in time and defence costs are significant (Rhoades 1979; Karban 2011). While these costs have likely been reduced through evolution by being selected against, the wide variety of defences found in both marine and terrestrial organisms suggests the presence of influential trade-offs to any beneficial defensive trait (Strauss *et al.* 2002; Agrawal 2011; Pančić *et al.* 2019; Grønning and Kiørboe submitted).

The failure of experiments to demonstrate costs may be due to the fact that experimental assessments have often been done under resource replete conditions, while costs may be more significant when resources are deficient (Karban 2011; Pančić & Kiørboe 2018; Kiørboe & Andersen 2019). The PST molecules are high in nitrogen with N:C ratio 4.6 times higher than average phytoplankton materials (Redfield 1958). Numerous studies have shown cell toxin content to be very low in nitrogen depleted cells (e.g. Boyer *et al.* 1987; Leong *et al.* 2004) even when exposed to a grazer threat (Selander *et al.* 2008). Trade-off costs may be trivial when nutrients and light are plentiful, but when available nitrogen is limiting and grazer biomass high, a fitness-optimization resource allocation model suggests a significant growth penalty to toxin production (Chakraborty *et al.* 2019).

Here, we quantify the benefits and costs of toxin production in *Alexandrium minutum* under different degrees of nitrogen limitation using both a chemostat approach and classical batch experiments. We induce grazer defences (toxin production) by adding grazer cues (copepodamides; Selander *et al.* 2015) and examine the growth rate response, the toxin production, and the efficiency of the defence by directly video recording the response of copepods to induced and non-induced cells. Following the predictions of the model of Chakraborty *et al.* (2019), we hypothesize that both induced and non-induced cells will be more toxic with increasing nitrogen availability, that the costs of increased toxicity of induced cells will be highest at intermediate nitrogen limitation, and that cells grown in excess of nitrogen will reap full benefit while paying negligible costs.

## Material and methods

### Phytoplankton

*Alexandrium minutum* strain GUMACC 83 were grown in B1 medium (Hansen 1989) at salinity 26, 18 °C, and an irradiance of *ca*. 150 μmol photons m^−2^ s^−1^ on a 12:12 light:dark cycle.

### Batch culture experiment

Six batch cultures of exponentially growing *A. minutum* were initiated at *ca*. 200 cells mL^−1^ in 2 L blue-cap glass flasks exposed to *ca*. 150 μmol photons m^−2^ s^−1^ on a 12:12 light:dark cycle and constant temperature of 18 °C. We used modified B1 medium with a nitrogen concentration of only 60 μM NO_3_^−^ to make sure that the cells eventually would be limited by nitrogen rather than other resources. The cultures were gently bubbled to avoid high pH limiting growth. pH was monitored using a PHM220 Lab pH meter (Radiometer Analytical, France). Three bottles were treated with copepodamides to induce increased toxin production (Selander *et al.* 2015) and three were used as controls. Copepodamides were extracted from freeze dried *Calanus finmarchius* (Calanus AS, Norway) according to (Selander *et al.* 2015) and exposed to the cultures by coating the inside of the bottle with a copepodamide mixture dissolved in methanol to a final concentration of 2 nM. Due to slow release and rapid degradation the average effective concentration is around 1−2% of the added concentration (Selander *et al.* 2019). The methanol was evaporated using N_2_ gas and the cultures transferred into the bottles after gentle mixing. This process was repeated and the culture transferred to a freshly coated flask every 24 hours during the treatment period to assure a continuous exposure to the cues (Selander *et al.* 2019). The controls received the same treatment but with methanol without copepodamides. Samples were taken every or every other day for cell abundance and nitrogen concentration while samples for toxin analysis and cellular carbon and nitrogen were taken at inoculation and then in tandem with the video experiments (see below) six times during the course of the experiment. Initial samples of cellular toxin-, carbon-, and nitrogen content were taken from the stock culture.

### Exponentially fed batch culture experiment

While nutrient concentration decline over time from high to limiting in the batch cultures, nutrients concentrations are near constant in a continuous cultures, thus allowing us to examine the effect of grazer induction at low, constant concentrations of nutrients. This may be important because the effect of grazer induction is time-lagged (Prevett *et al.* 2019). Dinoflagellates cannot tolerate vigorous mixing (Berdalet *et al.* 2007), and a classical chemostat cannot be used. Instead, we used exponentially fed batch cultures (EFB; Fischer *et al.* 2014). The EFB is similar to a chemostat except that there is no continuous outflow. The volume is instead reduced to initial value at each sampling occasion by removing medium and cells manually after gently mixing the culture. Growth medium is added continuously in a constant proportion of the increasing volume of the culture by exponentially increasing the inflow using a computer controllable multichannel peristaltic pump (IPC 16, Ismatec, Germany).

Six replicate cultures of *A. minutum* were set up as exponentially fed batch cultures in 1 L blue-cap glass bottles under the same conditions as above. Depending on the dilution rate, the initial culture volume varied between 250−500 mL. Four different dilution rates were used to vary cell growth rate: 0.05, 0.10, 0.20, and 0.40 d^−1^. The medium was prepared as B1 with reduced (80 μM) NO_3_^−^ in all the experiments except at the 0.10 d^−1^ dilution rate where the NO_3_^−^ concentration was 30 μM. The cultures were gently bubbled and pH was measured daily. At each dilution rate, the cultures were allowed up to ten days to achieve steady state before starting the experimental treatment. In some cases perfect steady-state was not achieved.

Copepodamides were exposed to the cultures daily as described above, using a nominal concentration of 0.63 nM. For the 0.2 d^−1^ dilution rate a second experiment was run with a copepodamide concentration of 6 nM. Samples for analysis of cell abundance and size, cell toxin content, cellular carbon and nitrogen, dissolved inorganic nitrogen, and copepod rejection rate were taken daily or every 2‒3 days during the 6‒10 day treatment period. Using the chemostat equations (Appendix 1) and assuming a maximum growth rate of 0.5 d^−1^ (Flynn *et al.* 1996) and a half saturation constant for nitrate of 0.5 μM (Brandenburg *et al.* 2018) the resulting nitrate concentration in the cultures should range from severe nitrogen limitation to saturation.

In the continuous cultures, the growth rate is fixed by the dilution rate and any growth rate response will materialize as a change in the steady state concentrations of cells and nutrients. Thus, if the cells respond to a cue by lowering their maximum growth or their affinity then in the chemostat at steady state the concentration of nutrients will increase, and the density of cells decrease in exposed compared to control cultures. The magnitude of the response can be computed from the chemostat equations (Appendix 1).

### Cell counts and cell size

Cell concentrations were determined by fixing a small volume of sample in acid Lugol’s solution to a final concentration of 1%. The entire chamber or at least 400 cells were counted per replicate in a Sedgewick-Rafter chamber using an inverted microscope (Olympus, Japan). 20 random cells from each sample were measured at 400× magnification (width-length) and cell volume was estimated by assuming a prolate spheroid shape (Hillebrand *et al.* 1999). Cell growth was calculated using the formula

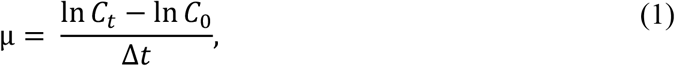

where *C_t_* is the cell concentration in units of cell number, biovolume, or cellular nitrogen per volume at the sampling occasion, *C_0_* is the concentration at the previous sampling occasion, and Δ*t* is the elapsed time in days. The dilution rate was added when calculating growth in the EFB experiments.

### Nitrate analysis

Subsamples for analysis of nitrate were filtered through a 0.2-μm syringe filter, and stored frozen at ‒20 °C until analysis. Nitrate was analysed by reduction to NO_x_ with VCl_3_ as the reducing reagent (Schnetger & Lehners 2014) on a Smartchem 200 (AMS Alliance, Italy). Concentrations below 0.5 μM were measured using an extended cuvette (100 mm, FireflySci, New York, USA) by UV-VIS spectrophometry.

### Toxin analysis

Samples (10−120 mL) for cellular toxin contents were filtered onto 25 mm Whatman GF/F glass fiber filters and frozen at ‒20 °C until extraction. 750 μL of 0.05 M acetic acid was added and samples were subjected to three cycles of freezing and thawing to lyse cells and extract toxins. The extracts were filtered through a GF/F filter and stored at ‒20 °C in glass HPLC vials until analysis. The samples from the batch experiment and the 0.2 d^−1^ dilution rate experiments (both low and high dose) were analysed with isocratic ion-pair chromatography followed by post column derivatization and fluorescent detection (Asp *et al.* 2004). We used a reversed phase C18 column (150×4 mm C18, 5 μm, Dr. Maisch GmBH, Germany). Samples from the 0.05, 0.10, and 0.40 d^−1^ dilution rate experiments were analyzed with mass spectrometric (MS/MS) detection on an Agilent 1200 HPLC system coupled to an Agilent 6470 triple quadruple detector with electrospray interface (Agilent Technologies, California, USA) equipped with a SeQuant zwitterionic hydrophilic interaction column (ZIC-HILIC, 2.1×150 mm, 5 μm, Merck, Germany) following the methods of (Turner & Tölgyesi 2019).

This particular strain of *A*. *minutum* is known to only produce gonyautoxins (GTX) 1‒4 (Franco *et al.* 1994; Selander *et al.* 2006). GTX standards 1‒4 were obtained from the National Research Council Canada (Halifax, Canada).

### Cellular carbon and nitrogen

Samples for cellular C and N were filtered onto pre-combusted (550 °C, 2 hours) 13 mm GF/C filters, packed in tin capsules and dried for 24 hours at 60 °C. The samples were kept dry at room temperature in a desiccator until analysis with a Thermo Scientific Flash 2000 Organic Elemental Analyzer (Thermo Scientific, Massachusetts, USA).

### Copepod feeding response

We directly observed individual copepod-cell interactions and recorded the fraction of captured cells that were rejected. We used the feeding-current feeding copepod *Temora longicornis* from a continuous culture that was maintained on a phytoplankton diet consisting of *Rhodamonas salina*, *Thalassioria weissflogii*, *Heterocapsa triquetra*, and *Oxyrrhis marina.*

Adult female copepods were tethered to a human hair by their dorsal surface using cyanoacrylate-based super glue (Xu *et al.* 2017). The tethered copepods were starved overnight in darkness at the same temperature (18 °C) and salinity as the cultures (26) before used for experiments. The tethered copepods are seemingly unaffected by the tethering and can live for many days while feeding, defecating, and producing eggs.

The feeding experiments took place in darkness. The untethered end of the hair was glued to a needle attached to a micromanipulator. The copepod was submerged in a 10×10×10 cm^3^ aquarium and *Alexandrium* cells were added to a final concentration of 100-200 cells mL^−1^. The experiment started when cells were added. A new copepod was used for each replicate culture. The water was gently mixed by a magnetic stirrer to keep the cells suspended. Three 10 minute sequences (0-10 minutes, 25-35 minutes, and 50-60 minutes) of 24 fps footage was recorded during a one hour period using a high speed camera (Phantom V210; Vision Research, New Jersey, USA) connected to a computer. The camera was equipped with lenses to get a field of view of 1.3 × 1 mm^2^. The video sequences were analysed to quantify prey capture, ingestion, and the fraction of cells that were rejected by the copepod (Xu *et al.* 2017).

### Statistical analysis

The effect of the copepodamide treatment in the batch experiment was analysed using a generalized additive mixed model (GAMM) in the ‘*gamm4*’ R package (Wood & Scheipl 2020). ‘Treatment’ and ‘Time’ were used as fixed effects and ‘Replicate’ as the random effect.

To analyse the effect of the copepodamide treatment in the chemostat experiments, we used a linear mixed effects model with ‘Time’, ‘Treatment’, and ‘Dilution rate’ as fixed effects, and ‘Replicate’ as the random effect, in the ‘*lmerTest’* R package (Kuznetsova *et al.* 2017). The analysis of the repeated ‘High’ copepodamide dose experiment was done separately. Significant differences between treatments were assessed through pairwise comparisons by estimated marginal means using the Satterthwaite degrees of freedom method. The random effect had a variance component that was close to zero when analysing some variables, but was retained in the model to incorporate the dependency of the response variable on the replications. Some variables were log-transformed to homogenize variances. Statistical tests were considered significant at the 0.05 level and are summarized in Appendix Tables S1, S2, S5 and S6.

## Results

Dinoflagellate concentrations and growth rates are in the following mainly reported in terms of cell number concentration (cells mL^−1^) and cell contents of C, N, and toxins on a per cell basis. Concentrations and growth rate in terms of cell volume concentration (μm^3^ mL^−1^) and the more sparsely monitored cellular nitrogen concentration (μg N mL^−1^) are reported in the online Appendix.

### Batch-culture experiment

Induced cultures grew faster than un-induced ones during the exponential phase and reached the stationary phase after around 14 days as the inorganic nitrogen in the cultures became depleted (Fig. 1A-C). The available nitrate in the culture medium was therefore used up at a higher rate in the induced treatment (Fig. 1C), because cell accumulation rate in terms of cellular nitrogen was also faster in induced than in un-induced cultures (Appendix 2 Fig. S1). Cellular nitrogen content and cell sizes initially increased and then decreased as nutrients were exhausted and growth ceased, but induced cells had a significantly higher nitrogen content (Fig. 1D) and were significantly smaller (Fig. 1G) than un-induced cells during the exponential growth phase before converging again with control values after 10 and 16 days, respectively. Cellular carbon content was significantly higher in induced than in un-induced cells (Fig. 1E). The differences in cellular C and N contents between induced and un-induced cultures are even more pronounced when normalized to cell volume (Appendix 2 Fig. S1). The carbon to nitrogen ratio increased over time as nitrate resources were exhausted but faster and to higher values in the induced treatment (Fig. 1F). Overall, cellular nitrogen content increased and cellular carbon content decreased with increasing growth rate and the contents of both nitrogen and carbon were higher in induced cells (Fig. 2, Appendix Table S2).

**Figure 1.**
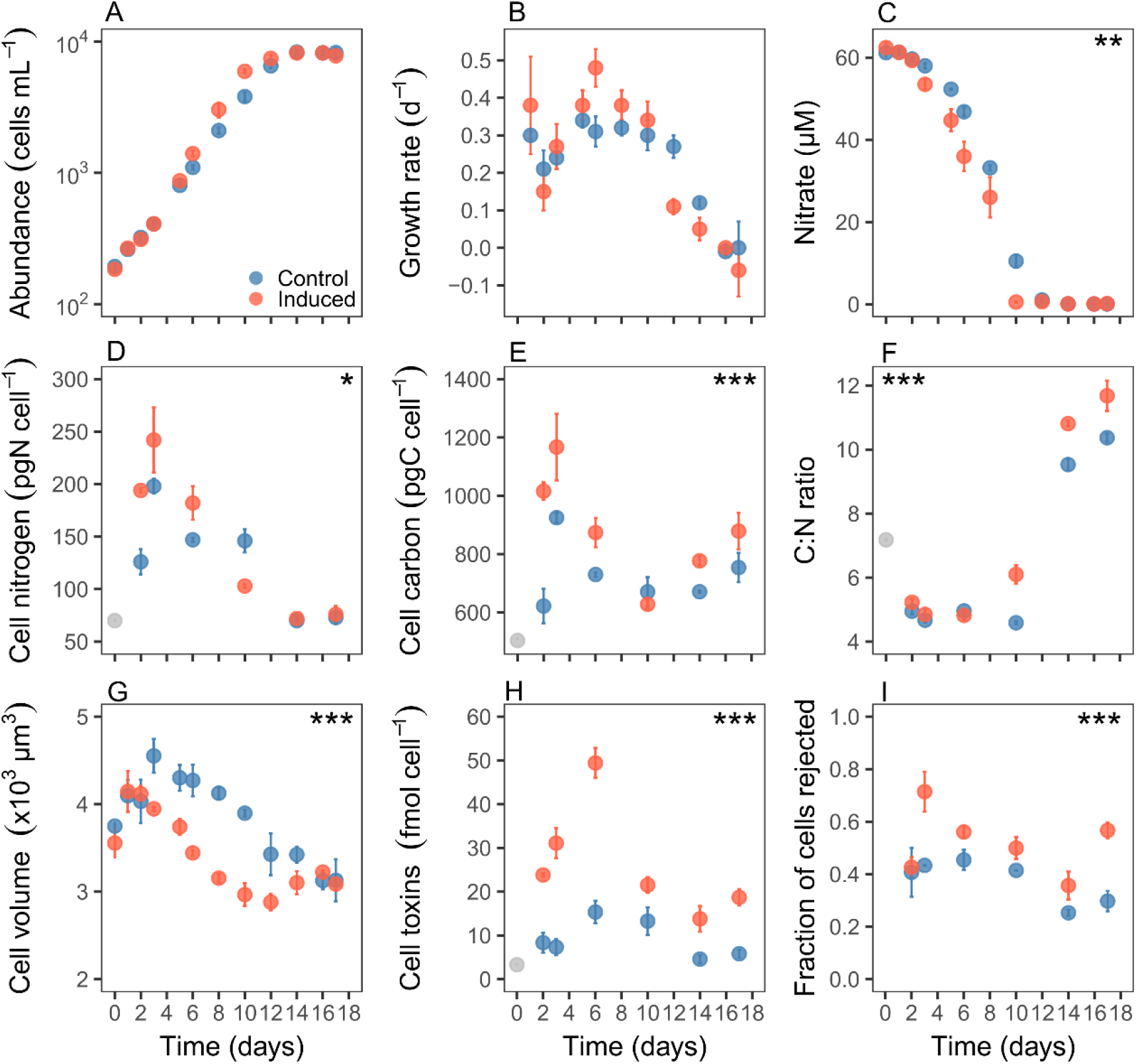
Change in (A) cell abundance (cells mL^−1^), (B) growth rate (d^−1^), (C) culture nitrate concentration (μM), (D) cell nitrogen content (pg N cell^−1^), (E) cell carbon content (pg C cell^−1^), (F) C:N ratio, (G) cell volume (μm^−3^), (H) cell toxin content (fmol cell^−1^), and (I) the fraction of cells rejected by copepods in the control (blue) and induced (red) *Alexandrium* cultures over time in the batch-culture experiment. The grey points in D, E, F, and H are initial values taken from the stock culture. Values are means and error bars are standard error (n = 3). Significant effects of the addition of copepodamides are indicated by the asterisks (*** *P* < 0.001, ** *P* < 0.01, * *P* <0.05).

**Figure 2.**
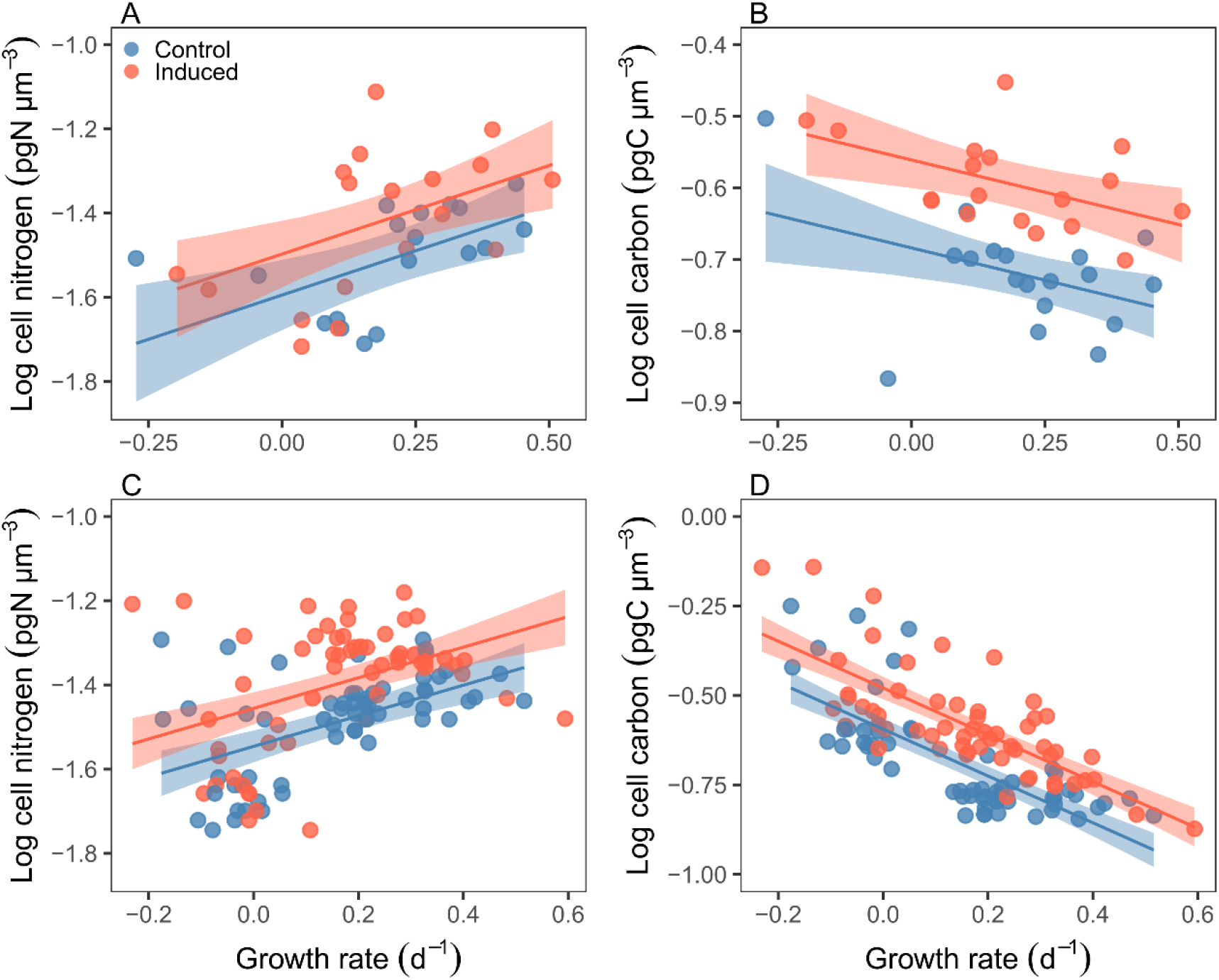
Relation between cell nitrogen (N, pg N μm^−3^) or cell carbon (C, pg C μm^−3^), and growth rate (GR, d^−1^) in the (A, B) batch and (C, D) EFB experiments. Growth rate is calculated from change in biovolume. Multiple linear regression was fit to the data: (A) control: log cell N = −1.594 + 0.420 × GR, induced: log cell N = −1.497 + 0.420 × GR (r^2^ = 0.27, *P* = 0.002); (B) control: log cell C = −0.684 − 0.181 × GR, induced: −0.561 − 0.181 × GR (r^2^ = 0.539, *P* < 0.001); (C) control: log cell N = −1.546 + 0.363 × GR, induced: −1.456 + 0.363 × GR (r^2^ = 0.28, *P* <0.001); and (D) control: log cell C = −0.594 – 0.654 × GR, induced: −0.479 – 0.654 × GR (r^2^ = 0.61, *P* < 0.001). The shaded areas are 95% confidence intervals.

Cell toxicity (320% increase relative to controls) peaked after six days of exposure in the induced treatments, after which it decreased throughout the rest of the exponential phase (Fig. 1H). Toxin production essentially reached zero after 14 days but cell toxicity remained stable at around 20 fmol cell^−1^ due to the low cell division rates. In the control treatment, cell toxicity followed the same temporal patterns but was throughout lower than in induced cells (Fig. 1H). Finally, a significantly higher fraction of induced than un-induced cells were rejected by copepods, demonstrating a clear benefit (Fig. 1I).

### Exponentially fed batch culture experiment

Cells in the 0.05, 0.10, and 0.40 d^−1^ dilution rate experiments grew at lower rates than the dilution rate and cell concentrations thus decreased over time (Fig. 3). It was only in the two 0.20 d^−1^ dilution rate experiments that the cells were able to keep up with the dilution rate (Fig. 3C). However, cell concentrations expressed in terms of cellular nitrogen per culture volume were near constant over times at the three lowest dilution rates, and the growth rates were similar to dilution rate except at the lowest and highest dilution rates (Figure 4D, Appendix 2 Fig. S2). Cell concentrations in nitrogen units were similar between induced and un-induced cultures at the highest dilution rate, but were generally higher in induced vs un-induced cultures at the lower dilution rates, suggesting higher growth rate or affinities of induced cells at limiting nutrient concentrations. The difference was significant only at the lowest dilution rate. Thus, if anything induced cells grow slightly faster, not slower, than un-induced cells at limiting nitrogen concentrations, consistent with the result of the batch experiment. The small differences in cell concentrations and the low sensitivity of estimates of affinity and maximum growth rate parameters to changes in cell concentration at low dilution rates makes the estimation of these parameters meaningless (Appendix 1).

**Figure 3.**
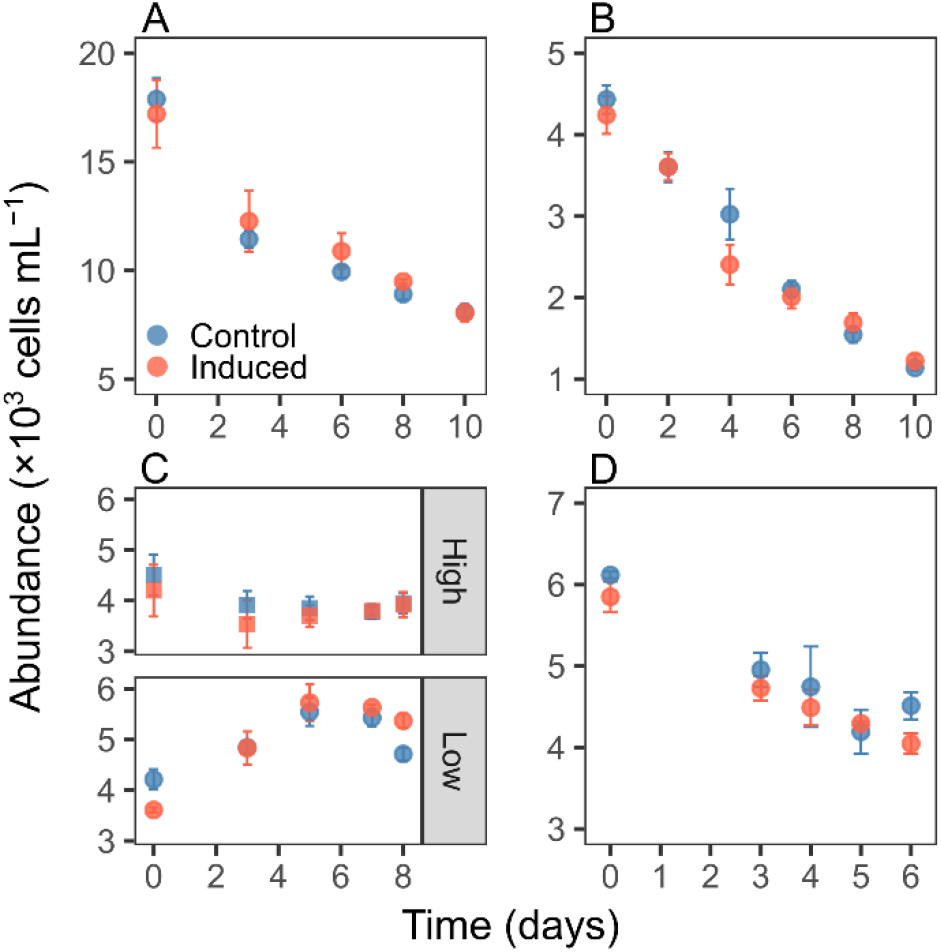
Change in cell abundance (cells mL^−1^) in the EFB at the different dilution rates. (A) 0.05 d^−1^, (B) 0.10 d^−1^, (C) 0.20 d^−1^ with high (6 nM) and low (0.63 nM) dose of copepodamides, (D) 0.40 d^−1^. The values are means and error bars show standard error (n = 3). Note the different y-axes scales.

**Figure 4.**
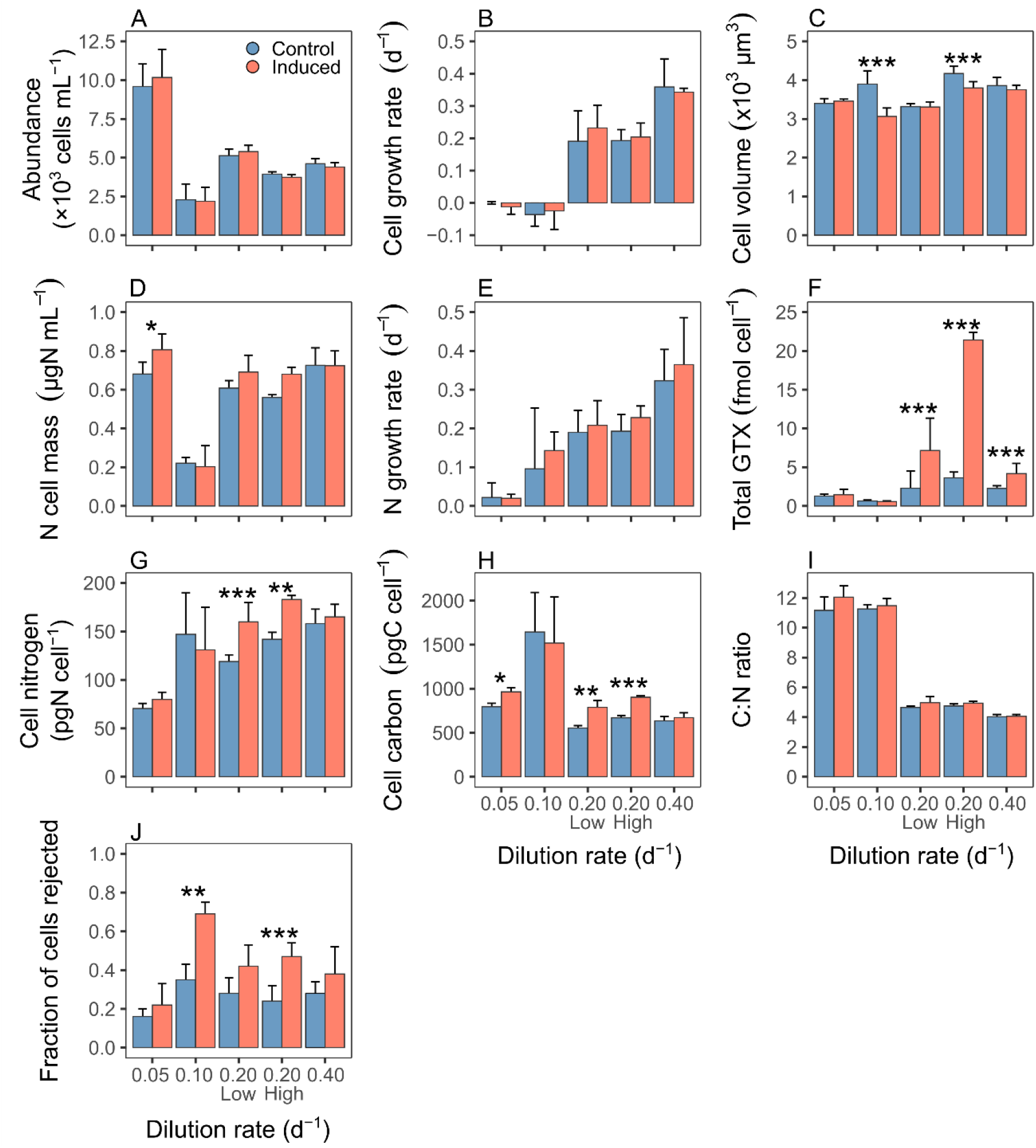
Summary of (A) cell abundance (×10^3^ cells mL^−1^), (B) cell growth rate (d^−1^), (C) cell volume (×10^3^ μm^3^), (D) nitrogen cell mass (μg N mL^−1^), (E) nitrogen growth rate (d^−1^), (F) cell toxin content (fmol cell^−1^), (G) cell nitrogen (pg N cell^−1^), (H) cell carbon (pg C cell^−1^), (I) cell C:N ratio, and (J) the fraction of cells rejected by copepods in the EFB experiments. Values are averaged over time during the treatment period and error bars show standard deviation (n = 4 in 0.05, 0.20, 0.40 d^−1^; n = 5 in 0.10 d^−1^; n = 3 in 0.10 d^−1^ C/N measurements). Significant differences between treatments within each dilution rate are indicated by asterisks (*** *P* < 0.001, ** *P* < 0.01, * *P* < 0.05).

Induced cells were significantly smaller than un-induced cells at intermediate dilution rates but similar at the lowest and highest rates, consistent with the pattern in the batch experiment. (Figure 4C). Cellular carbon increased and nitrogen contents decreased with growth rate and both were significantly higher in induced than un-induced cells, particularly at intermediate dilution rates (Figure 4G, H, Fig. 2). Cellular C:N ratio varied inversely with dilution rate and were slightly higher in induced than un-induced cells, all again consistent with the patterns in the batch experiment (Figure 4I, Fig. 2). As in the batch experiment, effects were more pronounced when expressed on a per volume basis (Appendix 2 Table S3, S4). The effect of varying the copepodamide dose from low (0.63 nM) to high (6 nM) in the 0.2 d^−1^ dilution rate experiment had a significant effect on (reduced) cell volume and also further increased toxin content relative to the controls (Fig. 4C, F).

Cells increased their toxicity per cell in all but the 0.05 and 0.10 d^−1^ experiments in response to the copepodamides (Fig. 4F) (The low toxicity in the 0.10 d^−1^ experiment is inconsistent with the high cellular nitrogen content and the high cell rejection rates; we suspect that the analysis is flawed). Consequently, the copepods generally rejected a larger fraction of induced cells, except at the lowest dilution rate (Fig. 4J; Appendix 2 Table S5). Overall, the fraction of rejected cells increased with increasing toxin content but the effect saturates at a rather low toxin content of ca. 20 fmol GTX cell^−1^ (Fig. 5).

**Figure 5.**
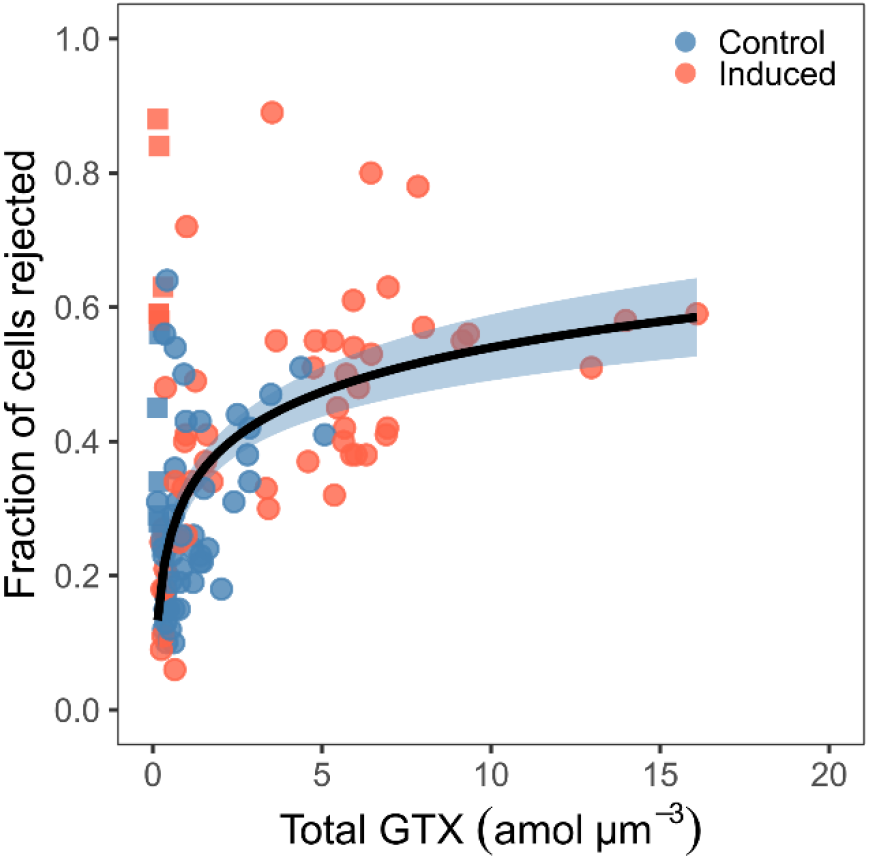
Relation between copepod rejection rate and cell toxin content normalized by volume (amol μm^−3^) in control (blue) and induced (red) cultures. Data are from both batch culture and EFB experiments. Squares are data from the 0.10 d^−1^ EFB experiment that are not included in the regression analysis. The regression line is Rejection = 0.320 + 0.219 × log GTX (r^2^ = 0.42, *P* < 0.001). The shaded area is the 95% confidence interval.

## Discussion

We set out to quantify the costs and benefits of toxin production in a dinoflagellate by comparing the performance of cells induced to express their defence with those that were not, and under different degrees of nitrogen limitation. Our experiments produced increasing nitrogen limitation with decreasing growth rates, in both batch and continuous cultures, as demonstrated by the relations between cellular nitrogen contents of the cells and their growth rate (Fig. 2A, C). We have utilized that toxin production in many dinoflagellates, including *A. minutum*, can be induced by grazer cues (Selander *et al.* 2006, 2015), thus allowing us to examine the same strain under different conditions. This is important because different strains of the same species may differ in many traits, including in their toxin profiles (e.g. Franco *et al.* 1994; Wohlrab *et al.* 2017). We note also that we examine the ‘private good’ grazer deterrent effect of the defence at the level of the individual. That is, we quantify the benefits that the individual cell that produces the toxin may enjoy. This is different from any toxic effects on the copepod that reduces its ability to grazer on further cells, which is a ‘public good’ (Driscoll *et al.* 2015). We had predicted that both benefits and costs would be small in nutrient starved cells, that benefits would be large but costs relatively small in nutrient replete cells, and that both benefits and costs would be high at intermediate nitrogen levels.

### Defence trade-offs

The benefits of toxin production were clear and largely followed the pattern predicted, and the results were consistent between the two types of experiments: induced cells have up to 3 times higher chance of being rejected by the copepod than un-induced cells, and the chance of rejection was directly related to the toxin content of the cells. This confirms previous reports of reduced grazing on induced dinoflagellates (Selander *et al.* 2006), but the demonstration at the individual cell level is novel. It is well established that nitrogen-starved cells produce no or very little toxins (Flynn *et al.* 1994; Selander *et al.* 2008) and, hence, enjoy little or no defence from toxins. In the batch experiment, cells remained toxic even in the stationary phase due to low cell division rate, but they did not produce new toxins. Even in nutrient replete conditions where cells continue to divide, it may take up to 14 days for induced *A. tamarense* to return to control levels after being removed from exposure to copepod cues (Selander *et al.* 2012).

The efficiency of the defence was not as high as that reported for other strains and species of *Alexandrium* spp. Thus, Xu & Kiørboe (2018) found that more than 90% of the cells of some toxic *Alexandrium* species/strains were rejected by a copepod, but also that other strains containing toxins were readably consumed by the copepod. Neither Xu *et al.* (2017) nor Teegarden *et al.* (2008) were consequently able to relate the efficiency of the defence to the composition and concentration of specific toxins in the cells of *Alexandrium* spp. Here, we have established a direct relation between the cells’ content of the GTX toxin and the efficiency of the defence. Transcriptional studies of grazer induced diatoms (Amato *et al.* 2018) reveals a rather massive response to grazer presence, with hundreds of genes being up-, or down-regulated. Thus is it quite possible that rejection rate resulted from additional up-regulated traits, acting in concert with the toxins.

It has been notoriously difficult to demonstrate the cost part of defence trade-offs in phytoplankton (Pančić & Kiørboe 2018), and this study is no exception. Ideally, ‘costs’ should be quantified in terms of reduced cell division rate. We found no reduction in the growth rate nor in nutrient affinity of the cells, even at nutrient deplete conditions.

However, we document a number of very clear effects of induction in addition to enhanced toxin production, i.e., elevated cellular contents of C and N, a reduction in cell size, and even an increase in cell division rate, with the effects being most pronounced at intermediate nutrient levels. The responses are consistent between the batch and the continuous cultures. However, it is not obvious whether these responses can be considered ‘costs’ or as parts of the defence.

The expectation of reduced cell division rate of grazer-induced, nitrogen-limited cells is based on the nitrogen requirements for PST production in *Alexandrium tamarense* as worked out by (Chakraborty *et al.* 2019). However, we had overlooked that the *A. tamarense* they considered produces high amounts of toxins, equivalent to the ~10^−9^ μg N μm^−3^, compared to the 1-2 orders of magnitude lower contents of ~10^−10^−10^−11^ μg N μm^−3^ in *A. minutum* found during our study. Thus, the biochemical syntheses costs and the N requirements for toxins are minute in this species. Thus, the higher N-uptake and -content of induced cells cannot be explained by direct requirements for nitrogen and energy for toxin production.

It is well established that cells shrink in size when nutrient limited (Peter & Sommer 2015; Garcia *et al.* 2016), as also seen most clearly in the batch experiment (Fig. 1G). However, and more important, induction by grazer cues causes cells to shrink, by up to 25% in volume relative to un-induced cells. This has two implications. First, a smaller cell volume implies a higher *concentration* of toxins in the cells. It is reasonable to assume that the copepods respond to the concentration rather than the contents of toxins, and the shrinking of the cells may therefore be adaptive and part of the defence. A similar consistent response in cell size to grazer cues has been found in seven species of diatoms (Grønning & Kiørboe submitted). For the diatoms, smaller cell sizes implies higher concentrations of biogenic silica and, therefore, a stronger protective shell that makes the cells less susceptible to copepod grazing (Pančić *et al.* 2019).

The second potential implication of cell shrinking is a higher affinity for dissolved nutrients. To first order, affinity scales inversely with cell radius due to the nature of molecular diffusion (Kiørboe 1993) and it is well established experimentally that the volume-specific nutrient uptake indeed increases with decreasing cell size (Edwards *et al.* 2012; Lindemann *et al.* 2016). Thus, a 25% decrease in cell volume, corresponds to an 8 % decrease in radius and a corresponding increase in affinity. This, in fact, may account for the elevated nitrogen uptake, nitrogen content, and growth rate of induced cells when cells start being nutrient limited, as most clearly seen in the batch experiment (Fig. 1). If the decrease in cell size is an adaptation to increase toxin concentration, then the elevated nitrogen assimilation and growth rate of induced cells is a secondary and maybe beneficiary effect.

The increased cell content found in induced cells may be due to thickening of their thecal plates, providing them with another possible defence. It is unclear if this has an effect on the copepods, but it has been shown that diatoms that increase their silica shell thickness experience reduced grazing from both juvenile and adult copepods (Pančić *et al.* 2019).

### Ecological and indirect costs of defence

In addition to direct costs, defences may also come with indirect ecological costs that do not manifest in simplified laboratory settings (Strauss *et al.* 2002). This includes, e.g., increased sinking rate or reduced swimming speed that may inflict fitness costs in nature (Lürling & Van Donk 2000; Selander *et al.* 2011). A possible ecological cost of the reduced cell size recorded here is elevated predation risk. In general, mortality rate of plankton organisms scale inversely with their volume to power 0.25 (Kiørboe 2008), and a 25% decrease in volume thus implies a 7.5% increase in predation mortality from other predators than copepods. Copepods and other larger herbivores are probably the most important grazers on dinoflagellate, thus the more than 50% decrease in copepod grazing pressure of induced cells more than outweighs the cost in most situations, and toxin production increases the fitness of the cell.

Theory of inducible defences predicts that defences should only be inducible if they are associated with a cost (Tollrian & Harvell 1999). The number of studies unable to detect costs associated with induced defences in phytoplankton suggests that additional factors may be at play. Recent advancements in genome sequencing reveals that a substantial part of the genome may be dedicated to secondary metabolism, up to one fifth in some cyanobacteria (Leao *et al.* 2017). Keeping a single biosynthetic pathway active may inflict a very limited cost whereas the cost for constant activation of one fifth of the genome will be substantial. Thus the evolution of inducible defences may be driven not by the allocation of resources to a single pathway, but the necessity to avoid allocation to all defence systems simultaneously. This is but a corollary hypothesis to the optimal defence theory, but one that may explain the lack of detectable costs in some induced responses to herbivory.

In conclusion, we found a complex nutrient-dependent response of a dinoflagellate to copepod cues: increase toxicity with implied lower predation risk, higher cellular contents of carbon and nitrogen, reduced cell size, and higher growth rate. Most of these responses may be beneficial to the cells, while we found no strong indication of costs. Because dinoflagellates are not Darwinian demons, the necessary costs are most likely indirect or ecological that are apparent only in nature.

## Acknowledgements

We thank Benni Winding Hansen for CHN measurements, Jack Melbye for maintaining copepod cultures, Colin Stedmon for assistance with NO_3_^−^ measurements, Josephine Grønning for copepodamide extractions, Aurore Maureaud for providing the script for the mixed model, Daniël van Denderen for providing statistical assistance, and Per Juel Hansen for rewarding discussions.

