## Appendix 1 for "Costs and benefits of toxin production in a dinoflagellate"

The chemostat equation assuming a Michaelis-Menten functional response in nutrient uptake to nutrient concentration reads

$$\frac{dB}{dt} = \mu_{max}\left(\frac{\alpha N(t)}{\mu_{max} + \alpha N(t)}\right)B - DB \tag{1}$$

$$\frac{dN}{dt} = -\mu_{max}\left(\frac{\alpha N(t)}{\mu_{max} + \alpha N(t)}\right)B + D\big(N_i - N(t)\big), \tag{2}$$

where $B$ is the phytoplankton biomass in culture (as cellular mol of nitrogen vol$^{-1}$), $D$ is the dilution rate (time$^{-1}$), $\mu_{max}$ the maximum growth rate (time$^{-1}$), $\alpha$ the affinity for nitrate-nitrogen (vol cellular mol of nitrogen$^{-1}$ time$^{-1}$), $N$ the nutrient concentration in culture (mol N vol$^{-1}$), and $N_i$ the nutrient concentration in inflow water (mass vol$^{-1}$). Solving for steady state yields

$$\hat{N} = \frac{D\mu_{max}}{\alpha(\mu_{max} - D)} \tag{3}$$

$$\hat{B} = N_i - \frac{D\mu_{max}}{\alpha(\mu_{max} - D)} = N_i - \hat{N}. \tag{4}$$

While eq. (3) and (4) are two equations with two unknowns ($\mu_{max}$, $\alpha$), they are not independent and one cannot find both unknowns. With one fixed, one can compute the other.

If the cells respond to a cue by lowering their maximum growth rate and/or its affinity then in the chemostat at steady state the concentration of nutrient will increase, and the density of cells decrease. Assuming first that the response is solely in the maximum growth rate, we can compute the response from the new steady state concentrations, dilution rate, and (known) affinity by solving either eq. (3) or eq. (4) for $\mu_{max}$:

$$\mu_{max} = \frac{\hat{N}\alpha D}{\hat{N}\alpha - D} \text{ or } \mu_{max} = \frac{D\alpha(N_i - \hat{B})}{\alpha(N_i - \hat{B}) - D}. \tag{5}$$

Alternatively, assume that the cells respond by lowering their affinity, then we can similarly compute the response in affinity from the steady state concentrations, the – known or assumed – maximum growth rate, and dilution rate:

$$\alpha = \frac{D\mu_{max}}{\hat{N}(\mu_{max} - D)} \text{ or } \alpha = \frac{-D\mu_{max}}{\hat{B}(\mu_{max} - D) - N_i(\mu_{max} + D)}. \quad (6)$$

Due to the shape of the functional response, these equations do not provide accurate absolute estimates, but do provide estimates of relative changes in maximum growth rate or affinity. However, and unfortunately, for low dilution rates, where according to our hypothesis changes in the parameters in response to grazer cues is expected to be largest, steady state cell concentrations are not very sensitive to changes in the parameters.
