## Appendix 2 for "Costs and benefits of toxin production in a dinoflagellate"

Appendix 2 Table S1. Summary statistics for the generalized additive mixed model (GAMM) used to assess the effect of the copepodamide treatment in the batch experiment.

| | | Estimate | Std. error | t value | *P* |
|---|---|---|---|---|---|
| Cell volume | Intercept | 3844.59 | 40.24 | 95.54 | <0.001 |
| | Treatment | −397.48 | 56.91 | −6.99 | <0.001 |
| Nitrate | Intercept | 31.98 | 0.72 | 44.22 | <0.001 |
| | Treatment | −3.17 | 1.02 | −3.11 | 0.003 |
| Cell nitrogen | Intercept | 118.68 | 4.54 | 26.13 | <0.001 |
| | Treatment | 15.57 | 6.42 | 2.42 | 0.022 |
| Cell carbon | Intercept | 696.74 | 19.93 | 34.96 | <0.001 |
| | Treatment | 138.36 | 28.18 | 4.91 | <0.001 |
| C:N ratio | Intercept | 6.61 | 0.07 | 89.99 | <0.001 |
| | Treatment | 0.63 | 0.10 | 6.08 | <0.001 |
| Cell toxin | Intercept | 8.28 | 0.98 | 8.49 | <0.001 |
| | Treatment | 14.81 | 1.38 | 10.73 | <0.001 |
| Rejection | Intercept | 0.38 | 0.03 | 13.82 | <0.001 |
| | Treatment | 0.15 | 0.04 | 3.89 | <0.001 |

Appendix 2 Table S2. Summary statistics for multiple regression analysis on the relationship between cell nitrogen (pg N μm$^{-3}$) or carbon (pg C μm$^{-3}$), and growth (d$^{-1}$) in the batch and exponentially fed batch culture (EFB) experiments. Growth rate is calculated from change in biovolume (μm$^{3}$ mL$^{-1}$). 'Lower' and 'Upper' refers to 95% confidence intervals.

| | | | Estimate | Std. Error | Lower | Upper | *P* |
|---|---|---|---|---|---|---|---|
| Batch | Log cell nitrogen | Intercept | −1.594 | 0.041 | −1.677 | −1.512 | <0.001 |
| | | Growth | 0.420 | 0.126 | 0.164 | 0.678 | 0.002 |
| | | Treatment | 0.097 | 0.044 | 0.008 | 0.187 | 0.034 |
| | Log cell carbon | Intercept | −0.684 | 0.020 | −0.725 | −0.643 | <0.001 |
| | | Growth | −0.181 | 0.062 | −0.307 | −0.054 | 0.007 |
| | | Treatment | 0.123 | 0.022 | 0.078 | 0.167 | <0.001 |
| EFB | Log cell nitrogen | Intercept | −1.546 | 0.019 | −1.584 | −1.508 | <0.001 |
| | | Growth | 0.363 | 0.067 | 0.230 | 0.497 | <0.001 |
| | | Treatment | 0.090 | 0.022 | 0.045 | 0.135 | <0.001 |
| | Log cell carbon | Intercept | −0.594 | 0.016 | −0.625 | −0.563 | <0.001 |
| | | Growth | −0.654 | 0.055 | −0.764 | −0.545 | <0.001 |
| | | Treatment | 0.115 | 0.017 | 0.078 | 0.152 | <0.001 |

Appendix 2 Table S3. Summary for exponentially fed batch culture experiments. The values are averaged over time and show means ± standard deviation.

| D. rate | Treatment | Dose | Abundance | Cell growth | N cell mass | N growth | Cell volume | Cell N | Cell C | C:N ratio | Cell toxin | Rejection |
|---|---|---|---|---|---|---|---|---|---|---|---|---|
| $d^{-1}$ | | | cells $mL^{-1}$ | $d^{-1}$ | μg N $mL^{-1}$ | $d^{-1}$ | $\mu m^{-3}$ | pg N $cell^{-1}$ | pg C $cell^{-1}$ | | fmol $cell^{-1}$ | |
| 0.05 | Control | Low | 9591±1450 | 0.000±0.004 | 0.681±0.061 | 0.022±0.038 | 3399±126 | 70.5±5.25 | 795±40 | 11.16±0.91 | 1.28±0.23 | 0.16±0.04 |
| 0.05 | Induced | Low | 10173±1816 | -0.012±0.023 | 0.806±0.082 | 0.020±0.010 | 3459±51 | 79.8±7.48 | 956±47 | 12.05±0.76 | 1.45±0.69 | 0.22±0.11 |
| 0.10 | Control | Low | 2282±1022 | -0.036±0.036 | 0.221±0.029 | 0.096±0.157 | 3899±343 | 147±43 | 1644±448 | 11.26±0.29 | 0.63±0.14 | 0.35±0.08 |
| 0.10 | Induced | Low | 2187±904 | -0.025±0.057 | 0.203±0.108 | 0.143±0.048 | 3065±217 | 131±44 | 1518±524 | 11.48±0.49 | 0.58±0.08 | 0.69±0.06 |
| 0.20 | Control | Low | 5133±416 | 0.191±0.094 | 0.608±0.039 | 0.190±0.057 | 3317±78 | 119±6.9 | 553±28 | 4.65±0.10 | 2.29±2.22 | 0.28±0.08 |
| 0.20 | Induced | Low | 5391±402 | 0.232±0.070 | 0.691±0.087 | 0.208±0.064 | 3308±138 | 160±20 | 789±79 | 4.97±0.41 | 7.16±4.18 | 0.42±0.11 |
| 0.20 | Control | High | 3942±145 | 0.193±0.034 | 0.560±0.014 | 0.193±0.043 | 4173±190 | 142±7.1 | 669±27 | 4.75±0.14 | 3.62±0.76 | 0.24±0.08 |
| 0.20 | Induced | High | 3738±163 | 0.204±0.043 | 0.680±0.036 | 0.228±0.030 | 3800±160 | 183±4.1 | 903±17 | 4.94±0.13 | 21.40±0.90 | 0.47±0.08 |
| 0.40 | Control | Low | 4611±333 | 0.359±0.087 | 0.726±0.090 | 0.323±0.081 | 3862±211 | 158±15 | 635±49 | 4.03±0.15 | 22.7±0.34 | 0.28±0.06 |
| 0.04 | Induced | Low | 4390±289 | 0.343±0.012 | 0.724±0.074 | 0.365±0.121 | 3753±119 | 165±13 | 670±57 | 4.06±0.12 | 4.18±1.31 | 0.38±0.14 |

Appendix 2 Table S4. Summary for exponentially fed batch culture experiments, normalized by cell volume to account for differences in size. The values are averaged over time and show ± standard deviation.

| Dil. rate $d^{-1}$ | Treatment | Dose | Biovolume $\times 10^7$ $\mu m^3$ $mL^{-1}$ | Growth $d^{-1}$ | Cell N pg N $\mu m^{-3}$ | Cell C pg C $\mu m^{-3}$ | Cell toxin amol $\mu m^{-3}$ |
|---|---|---|---|---|---|---|---|
| 0.05 | Control | Low | 3.269±0.562 | -0.011±0.024 | 0.021±0.002 | 0.234±0.016 | 0.40±0.06 |
| 0.05 | Induced | Low | 3.520±0.658 | -0.015±0.030 | 0.023±0.002 | 0.278±0.015 | 0.45±0.20 |
| 0.10 | Control | Low | 0.870±0.340 | -0.052±0.076 | 0.036±0.010 | 0.40±0.11 | 0.16±0.03 |
| 0.10 | Induced | Low | 0.660±0.256 | -0.066±0.120 | 0.042±0.015 | 0.48±0.17 | 0.19±0.03 |
| 0.20 | Control | Low | 1.705±0.113 | 0.217±0.042 | 0.040±0.003 | 0.167±0.01 | 0.69±0.68 |
| 0.20 | Induced | Low | 1.783±0.166 | 0.234±0.038 | 0.053±0.007 | 0.261±0.03 | 2.23±1.37 |
| 0.20 | Control | High | 1.644±0.048 | 0.208±0.024 | 0.034±0.001 | 0.160±0.01 | 0.88±0.16 |
| 0.20 | Induced | High | 1.421±0.111 | 0.203±0.064 | 0.048±0.003 | 0.238±0.01 | 5.40±0.38 |
| 0.40 | Control | Low | 1.772±0.066 | 0.372±0.043 | 0.041±0.005 | 0.165±0.01 | 0.59±0.09 |
| 0.40 | Induced | Low | 1.651±0.141 | 0.340±0.053 | 0.044±0.004 | 0.180±0.02 | 1.11±0.35 |

Appendix 2 Table S5. Type III analysis of variance (ANOVA) on the fixed effects in the linear mixed models used to analyze the effect of the copepodamide treatment in low dose (0.63 nM) EFB experiments. *P*-values are provided via Satterthwaite's of freedom method. Some variables were log-transformed to homogenize variances. Only variables where the post-hoc test found significant differences between treatments are reported.

| | | Sum Sq. | NumDF | DenDF | F | *P* |
|---|---|---|---|---|---|---|
| Cell volume | Treatment | 53547 | 1 | 92 | 0.856 | 0.356 |
| | Time | 380975 | 1 | 75 | 6.117 | 0.016 |
| | Dil. rate | 2944857 | 3 | 16 | 15.762 | <0.001 |
| | Treatment×Time | 35618 | 1 | 75 | 0.572 | 0.452 |
| | Treatment×Dil rate | 2819502 | 3 | 16 | 15.091 | <0.001 |
| Nitrogen cell mass | Treatment | 0.00017 | 1 | 79 | 2.856 | 0.854 |
| | Time | 0.08307 | 1 | 65 | 16.868 | <0.001 |
| | Dil. rate | 0.96757 | 3 | 18 | 65.495 | <0.001 |
| | Treatment×Time | 0.00279 | 1 | 65 | 0.566 | 0.455 |
| | Treatment×Dil rate | 0.02323 | 3 | 18 | 1.573 | 0.231 |
| Cell toxins | Treatment | 161.76 | 1 | 91 | 44.823 | <0.001 |
| | Time | 33.64 | 1 | 74 | 9.321 | 0.003 |
| | Dil. rate | 1579.71 | 3 | 15 | 145.906 | <0.001 |
| | Treatment×Time | 7.71 | 1 | 75 | 2.136 | 0.148 |
| | Treatment×Dil rate | 909.56 | 3 | 15 | 84.009 | <0.001 |
| Log cell nitrogen* | Treatment | 0.00030 | 1 | 80 | 0.065 | 0.800 |
| | Time | 0.06384 | 1 | 80 | 13.609 | <0.001 |
| | Dil. rate | 1.54569 | 3 | 80 | 109.838 | <0.001 |
| | Treatment×Time | 0.00494 | 1 | 80 | 1.053 | 0.308 |
| | Treatment×Dil rate | 0.08440 | 3 | 80 | 5.997 | <0.001 |
| Log cell carbon* | Treatment | 0.00059 | 1 | 80 | 0.126 | 0.723 |
| | Time | 0.94938 | 1 | 80 | 10.573 | 0.002 |
| | Dil. rate | 1.08142 | 3 | 80 | 77.176 | <0.001 |
| | Treatment×Time | 0.01077 | 1 | 80 | 2.305 | 0.133 |
| | Treatment×Dil rate | 0.11553 | 3 | 80 | 8.245 | <0.001 |
| Log rejection* | Treatment | 0.15072 | 1 | 52 | 4.334 | 0.043 |
| | Time | 0.01625 | 1 | 52 | 0.467 | 0.497 |
| | Dil. rate | 1.69767 | 3 | 52 | 16.271 | <0.001 |
| | Treatment×Time | 0.03181 | 1 | 52 | 0.915 | 0.343 |

| | | | | | |
|---|---|---|---|---|---|
| Treatment×Dil rate | 0.09947 | 3 | 52 | 0.953 | 0.422 |

*: Random effect variances estimated as (close to) zero.

Appendix 2 Table S6. Type III analysis of variance (ANOVA) on the fixed effects in the linear mixed models used to analyze the effect of the copepodamide treatment in high dose (6 nM) repeated 0.2 $d^{-1}$ dilution rate EFB experiment. *P*-values are provided via Satterthwaite's degrees of freedom method. Only variables where the post-hoc test found significant differences between treatments are reported.

| | | Sum Sq. | NumDF | DenDF | F | *P* |
|---|---|---|---|---|---|---|
| Cell volume* | Treatment | 27104 | 1 | 20 | 0.890 | 0.357 |
| | Time | 431937 | 1 | 20 | 14.183 | 0.001 |
| | Treatment×Time | 17325 | 1 | 20 | 0.569 | 0.459 |
| Cell toxins | Treatment | 136.698 | 1 | 18 | 37.723 | <0.001 |
| | Time | 0.306 | 1 | 16 | 0.085 | 0.775 |
| | Treatment×Time | 0.536 | 1 | 16 | 0.148 | 0.706 |
| Cell nitrogen | Treatment | 2005 | 1 | 20 | 11.010 | 0.003 |
| | Time | 114 | 1 | 16 | 0.682 | 0.440 |
| | Treatment×Time | 219 | 1 | 16 | 1.212 | 0.287 |
| Cell carbon* | Treatment | 54997 | 1 | 20 | 14.033 | 0.001 |
| | Time | 224 | 1 | 20 | 0.057 | 0.814 |
| | Treatment×Time | 3094 | 1 | 20 | 0.789 | 0.385 |
| Log rejection* | Treatment | 0.164 | 1 | 17 | 9.460 | 0.007 |
| | Time | 0.001 | 1 | 17 | 0.010 | 0.921 |
| | Treatment×Time | 0.033 | 1 | 17 | 1.919 | 0.184 |

*: Random effect variances estimated as (close to) zero.

Appendix 2 Figure S1.

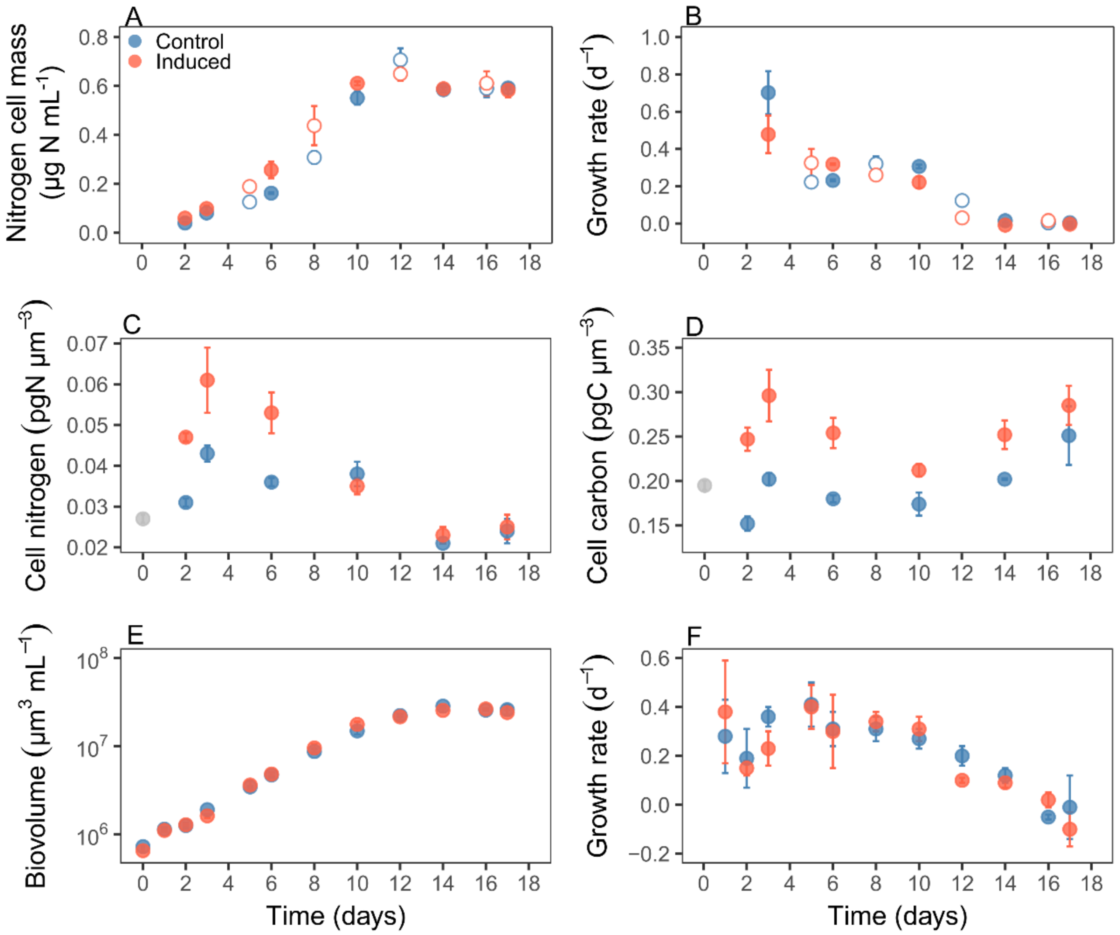


Figure S1. Change in (A) nitrogen cell mass (μg N mL$^{-1}$), (B) growth calculated from A (d$^{-1}$), (C) cell nitrogen (pg N μm$^{-3}$) and (D) carbon (pg C μm$^{-3}$) per cell volume, (E) biovolume (μm$^3$ mL$^{-1}$), and (F) growth (d$^{-1}$) calculated from E, over time in the batch culture experiment. The grey points in C and D are initial values taken from the stock culture. Empty symbols in A and B are interpolated values to improve resolution. Values are means and error bars show standard error (n = 3).

Appendix 2 Figure S2.

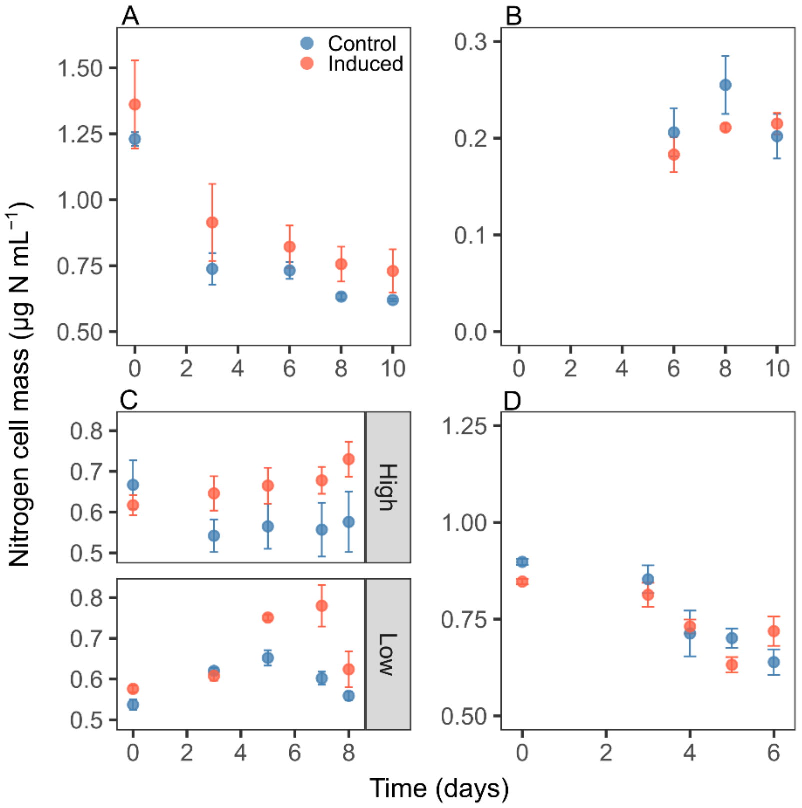


Figure S2. Change in nitrogen cell mass (µg N mL$^{-1}$) in the EFB at the different dilution rates. (A) 0.05 d$^{-1}$, (B) 0.10 d$^{-1}$, (C) 0.20 d$^{-1}$ with high (6 nM) and low (0.63 nM) dose of copepodamides, (D) 0.40 d$^{-1}$. The values are means and error bars show standard error (n=3). Note the different y-axes scales.
